# Ligand Gaussian accelerated molecular dynamics 2 (LiGaMD2): Improved calculations of ligand binding thermodynamics and kinetics with closed protein pocket

**DOI:** 10.1101/2022.12.16.520748

**Authors:** Jinan Wang, Yinglong Miao

## Abstract

Ligand binding thermodynamics and kinetics are critical parameters for drug design. However, it has proven challenging to efficiently predict ligand binding thermodynamics and kinetics from molecular simulations due to limited simulation timescales. Protein dynamics especially in the ligand binding pocket often plays an important role in ligand binding. Based on our previously developed Ligand Gaussian accelerated molecular dynamics (LiGaMD), here we present LiGaMD2 in which a selective boost potential was applied to both the ligand and protein residues in the binding pocket to improve sampling of ligand binding and dissociation. To validate the performance of LiGaMD2, the T4 lysozyme (T4L) mutants with open and closed pockets bound by different ligands were chosen as model systems. LiGaMD2 could efficiently capture repetitive ligand dissociation and binding within microsecond simulations of all T4L systems. The obtained ligand binding kinetic rates and free energies agreed well with available experimental values and previous modeling results. Therefore, LiGaMD2 provides an improved approach to sample opening of closed protein pockets for ligand dissociation and binding, thereby allowing for efficient calculations of ligand binding thermodynamics and kinetics.

## Introduction

ligand binding to target receptors plays a critical role in many fundamental biological processes^1^, as well as in the design of more effective and selective drugs for treating human diseases^2^. A number of experimental techniques^3^ have been developed to explore protein-small molecule interactions. For example, structural biology techniques^3b^ have been widely applied to determine protein-ligand complex structures. However, X-ray crystallography and cryo-electron microscopy (cryo-EM) could provide only static snapshots of protein-small molecule interactions. It is rather challenging for experimental methods to capture ligand binding and dissociation pathways and determine potential intermediate states of ligand binding to the protein target site.

Recently, ligand binding kinetics has been recognized to be critical for drug design^4^. The ligand dissociation rate (*k*_*off*_) or residence time (1/ *k*_*off*_) appears to correlate with drug efficacy better than ligand binding free energy. However, ligand binding kinetic rates have proven more challenging to predict, due to slow processes of ligand dissociation and rebinding^4b, 5^. With remarkable advances in computer hardware and method developments, conventional molecular dynamics (cMD) simulations nowadays are able to capture spontaneous ligand binding to target proteins and predict corresponding binding association rates (*k*_*on*_)^6^. However, it remains challenging to use cMD to simulate repetitive ligand binding and dissociation processes, precluding accurate prediction of ligand binding kinetic rates. For example, based on tens-of-microsecond cMD simulations, Shan et al. ^7^ successfully captured spontaneous binding of the Dasatinib drug to its target Src kinase and accurately predicted the ligand association rate (*k*_*on*_). However, no dissociation event was observed in the cMD simulations. Similar results were observed in the binding of benzene (BEN) to the L99A mutant of T4 lysozyme (T4L)^8^. The cMD simulations with lengths of 2 to 8 μs successfully captured BEN binding to the L99A T4L^8^. Tens to hundreds of repetitive guest binding and dissociation from the β-CD host were observed in microsecond cMD simulations^9^, which enabled accurate prediction of host-guest binding thermodynamics and kinetics. Tens-of-microsecond cMD simulations^6d^ captured repetitive binding and dissociation of six small-molecule fragments with weak millimolar binding affinities to the protein FKBP, which enabled accurate prediction of fragment binding free energies and kinetic rates. Nevertheless, cMD simulations have not captured repetitive binding and dissociation of typical small-molecule ligands to target proteins so far.

In this regard, enhanced sampling methods^10^ have been developed to extend accessible timescale of MD simulations, including Metadynamics^11^, Steered MD^12^, Umbrella Sampling^12a, 13^, Replica Exchange MD ^14^, Random Acceleration Molecular Dynamics (RAMD) ^15^, Scaled MD ^16^, accelerated MD (aMD) ^17^, Gaussian accelerated MD (GaMD) ^18^, Markov State Model (MSM)^8, 19^, Weighted Ensemble^20^, and so on. Using T4L as a model system, a total of ∼12 μs infrequent Metadynamics^11a, 11b^ simulations successfully captured 20 times of ligand binding and dissociation events and predicted the values of *k*_*on*_ and *k*_*off*_ at 3.5 ± 2 × 10^4^ M^−1^s^−1^ and 7 ± 2 s^−1^, being comparable to experimental values of 0.8±1×10^6^M^-1^s^-1^ and 950±20 s^-1^, respectively. However, Metadynamics requires predefined collective variables (CVs) before running the simulations. Thus, *a priori* knowledge of the systems is often required. In comparison, Weighted Ensemble^21^ and MSM^22^ combine a large number of short cMD simulations to predict ligand binding kinetic rates. For example, Weighted Ensemble^20b^ of a total of 29 μs cMD was able to accurately predict the dissociation rate of BEN from the L99A T4L as 1000s^-1^, being highly consistent with the experimental value of 950±200s^-1^. One MSM^8^ built on a total of 69 μs cMD simulations predicted the values of *k*_*on*_ and *k*_*off*_ at 21±91×10^6^ M^-1^s^-1^ and 311±130 s^-1^, being in reasonable agreement with experiment values of 0.8±1×10^6^M^-1^s^-1^ and 950±20 s^-1^, respectively. However, these calculations required very expensive computational resources.

GaMD was developed to provide both unconstrained enhanced sampling and free energy calculations of large biomolecules^18^. It works by applying a harmonic boost potential to reduce system energy barriers. The boost potential normally exhibits a near Gaussian distribution, which enables proper reweighting of the free energy profiles through cumulant expansion to the second order^18^. Ligand GaMD (LiGaMD)^23^ has recently been developed to allow for more efficient sampling of ligand dissociation and rebinding processes, being able to accurately predict ligand binding thermodynamics and kinetics. In LiGaMD, one selective boost was applied to the ligand non-bonded interaction potential energy to accelerate ligand dissociation. Another boost was applied on the remaining potential to facilitate ligand rebinding. Recently, an increasing number of studies suggested that the flexibility of proteins, particularly those with closed pockets, played an important role in ligand binding^8, 24^. Protein structural flexibility allows the opening, closing, and adaptation of the binding pocket, which are critical for ligand binding to protein^8, 25^. Additionally, enhanced sampling of the protein binding pocket has proven to significantly improve the efficiency and accuracy of simulating protein-ligand interaction^14d, 26^. Wang et al.^26^ proposed FEP/REST to improve the accuracy of ligand binding free energy calculation, which combined the free energy perturbation (FEP) and replica exchange with solute tempering (REST) to enhance sampling of protein residues in the binding site. FEP/REST was demonstrated to achieve much quicker simulation convergence and more accurate binding free energy calculations. Sugita et al.^14d^ developed the generalized replica exchange with solute tempering (gREST) to simulate small molecule binding to the L99A T4L. By enhanced sampling of ligand and protein residues in the binding site, the gREST simulations successfully captured ligand binding to the L99A T4L in a total of 2.4 μs simulations. The obtained free energy profiles of binders (BEN, ethylbenzene, and n-hexylbenzene) were distinct from those of nonbinders (phenol and benzaldehyde). Another study^25a^ from the same group combined the gREST with the replica exchange umbrella sampling (gREST/REUS) to capture the binding of compound PP1 to its target Src kinase. The gREST/REUS simulations^25a^ could capture multiple ligand binding and unbinding events.

Built on our previously developed LiGaMD, here we developed a novel approach termed LiGaMD2, in which a selective boost was added to both the ligand and protein pocket residues to accelerate ligand dissociation and binding. Various T4L mutants bound by BEN and indole (IND) with open and closed pockets were chosen as testing systems. The T4L mutants with small hydrophobic cavities that can accommodate different ligands have been widely used as model systems for benchmarking computational methods^27^. Microsecond LiGaMD2 simulations could capture repetitive ligand binding and unbinding processes in all the simulated T4L systems. LiGaMD2 thus enabled highly efficient and accurate prediction of ligand binding thermodynamics and kinetics. Simulation predictions agreed well with experimental binding free energy and kinetic rates. Since the chosen systems included different ligands and protein mutants with distinct binding pockets, the simulations validated the robustness of LiGaMD2. Furthermore, multiple ligand binding and dissociation pathways were identified by LiGaMD2 simulations, being highly consistent with published simulation results^8, 15a, 20b, 28^.

## Methods

### LiGaMD2: Selectively boosting both the ligand and protein pocket

LiGaMD is an enhanced sampling technique for characterizing ligand binding thermodynamics and kinetics. It works by adding a selective harmonic boost potential to the non-bonded ligand interaction potential energy. Detail of the LiGaMD method has been described in our previous study^29^. Here, we briefly describe the algorithm for further development of the LiGaMD2 method.

We consider a system of ligand *L* binding to a protein *P* in a biological environment *E*. The system comprises of *N* atoms with their coordinates 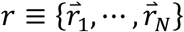 and momenta 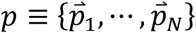. The system Hamiltonian can be expressed as:

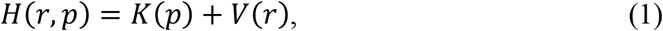

where *K* (*p*) and *V* (*r*) are the system kinetic and total potential energies, respectively. The protein P could be further divided into two parts: residues in the binding pocket (Pb) and other parts (Po). We decompose the potential energy into the following terms:

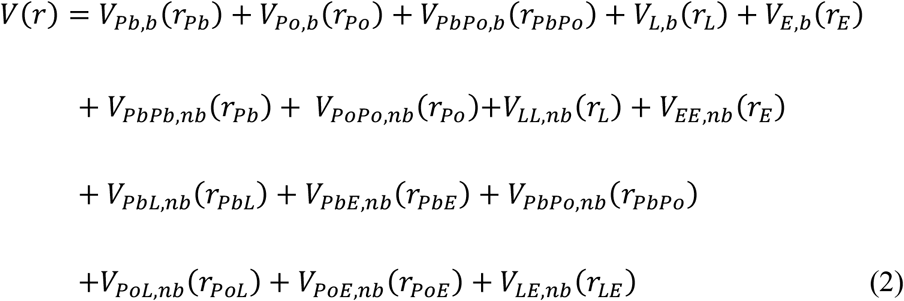

where *V*_*Pb,b*_, *V*_*Po,Pb*_, *V*_*L,b*_ and *V*_*E,b*_ are the bonded potential energies in the binding pocket of protein Pb, the remaining atoms of protein Po, ligand *L* and environment *E*, respectively. *V*_*PbPo, b*_(*r*_*PbPo*_) is the bonded potential energies involving atoms between the protein binding pocket and the other parts. *V*_*PbPb,nb*_, *V*_*PoPo,nb*_, *V*_*LL,nb*_ and *V*_*E,nb*_ are the self non-bonded potential energies in the protein binding pocket Pb, the remaining atoms of protein Po, ligand *L* and environment *E*, respectively. *V*_*PbPo,nb*_(*r*_*PbPo*_), *V*_*PbL,nb*_ (*r*_*PbL*_), *V*_*PbE,nb*_ (*r*_*PbEE*_), *V*_*PoL,nb*_ (*r*_*PoL*_), *V*_*PoE,nb*_ (*r*_*PoE*_) and *V*_*LE,nb*_are the non-bonded interaction energies between Pb-Po, Pb-L, Pb-E, Po-L, Po-E and L-E, respectively. According to classical molecular mechanics force fields^30^, the non-bonded potential energies are usually calculated as:

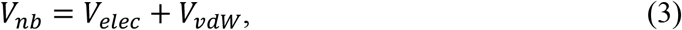

where *V*_*elec*_ and *V*_*vdw*_ are the system electrostatic and van der Waals potential energies. The bonded potential energies are usually calculated as

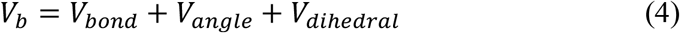

where *V*_*bond*_, *V*_*angle*_ and *V*_*dihedral*_ are the system bond, angle and dihedral potential energies. As mentioned above, flexibility of the protein pocket often plays a critical role in ligand binding. Therefore, the ligand essential potential energy in the LiGaMD2 is defined as

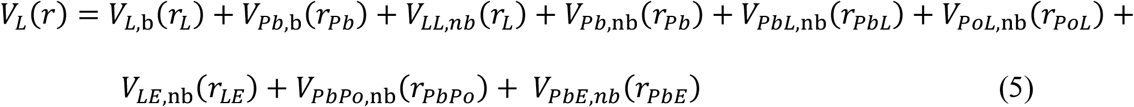

In the Pd-GaMD, we add boost potential selectively to the ligand essential potential energy according to the GaMD algorithm:

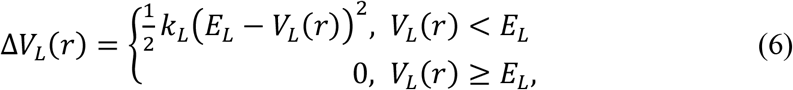

where *E*_*L*_ is the threshold energy for applying boost potential and *k*_*L*_ is the harmonic constant. The LiGaMD2 simulation parameters are derived similarly as in the previous LiGaMD. When *E* is set to the lower bound as the system maximum potential energy (*E=V*_*max*_), the effective harmonic force constant *k*_0_ can be calculated as:

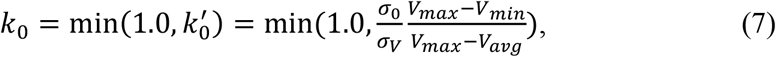

where *V*_*max*_, *V*_*min*_, *V*_*avg*_ and *σ*_*V*_ are the maximum, minimum, average and standard deviation of the boosted system potential energy, and *σ*_0_ is the user-specified upper limit of the standard deviation of Δ*V* (e.g., 10*k*_*B*_T) for proper reweighting. The harmonic constant is calculated as *k* = *k*_0_ · 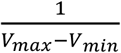 with 0 < *k*_0_ ≤ 1. Alternatively, when the threshold energy *E* is set to its upper bound 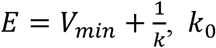 is set to:

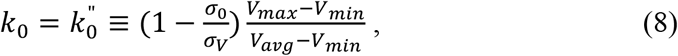

if 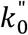 between 0 and 1. Otherwise, *k*_0_ is calculated using Eqn. (7).

In addition to selectively boosting the ligand and surrounding protein residues in the binding site, another boost potential is applied on the protein and solvent to enhance conformational sampling of the protein and facilitate ligand rebinding. The second boost potential is calculated using the total system potential energy other than the ligand essential potential energy as:

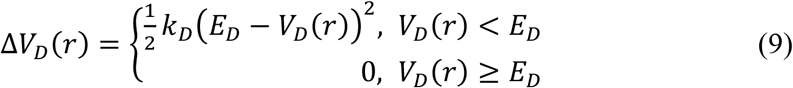

Where V_*D*_ is the total system potential energy other than the ligand essential potential energy, E_*D*_ is the corresponding threshold energy for applying the second boost potential and k_*D*_ is the harmonic constant. This leads to dual-boost LiGaMD2 with the total boost potential Δ*V*(*r*) = Δ*V*_*L*_ (*r*) + Δ*V*_*D*_ (*r*).

### Energetic Reweighting of LiGaMD2

To calculate potential of mean force (PMF)^31^ from LiGaMD2 simulations, the probability distribution along a reaction coordinate is written as *p*^*^(*A*). Given the boost potential 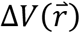 of each frame, *p*^*^(*A*) can be reweighted to recover the canonical ensemble distribution, *p* (*A*), as:

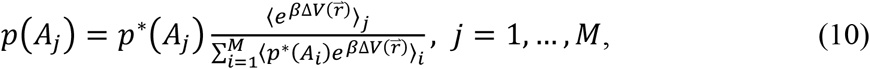

where *M* is the number of bins, *β* = *k*_*B*_*T* and 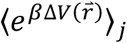 is the ensemble-averaged Boltzmann factor of 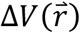 for simulation frames found in the *j*^th^ bin. The ensemble-averaged reweighting factor can be approximated using cumulant expansion:

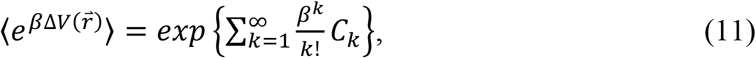

where the first two cumulants are given by

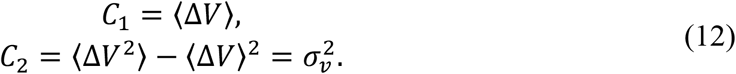

The boost potential obtained from LiGaMD2 simulations usually follows near-Gaussian distribution. Cumulant expansion to the second order thus provides a good approximation for computing the reweighting factor^18b, 32^. The reweighted free energy *F* (*A*) = −*k*_*B*_*T* ln *p* (*A*) is calculated as:

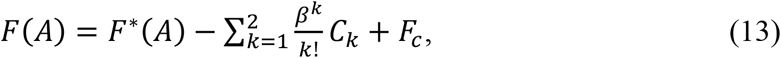

where *F*^*^(*A*) = −*k*_*B*_*T* ln *p*^*^(*A*) is the modified free energy obtained from LiGaMD2 simulation and *F*_*c*_ is a constant.

### Ligand binding free energy calculations from 3D potential of mean force

We calculate ligand binding free energy from 3D potential of mean force (PMF) of ligand displacements from the target protein as the following^33^:

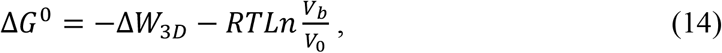

where *V*_0_ is the standard volume, *V*_*P*_ = ∫_*b*_ *e*^−*βW* (*r*)^ *dr* is the average sampled bound volume of the ligand with *β* = 1/*k*_*B*_*T, k*_*B*_ is the Boltzmann constant, *T* is the temperature, and Δ*W*_3*D*_ is the depth of the 3D PMF. Δ*W*_3*D*_ can be calculated by integrating Boltzmann distribution of the 3D PMF. *W* (*r*) over all system coordinates except the x, y, z of the ligand:

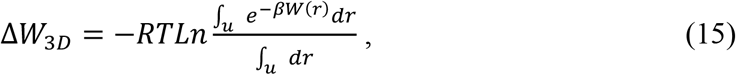

where *V*_*u*_ = ∫_*u*_*dr* is the sampled unbound volume of the ligand. The exact definitions of the bound and unbound volumes *V*_*P*_ and *V*_*u*_ are not important as the exponential average cut off contributions far away from the PMF minima^33b^. A python script “PyReweighting-3D.py” in the *PyReweighting* tool kit (http://miao.compbio.ku.edu/PyReweighting/)^34^ was applied for reweighting LiGaMD2 simulations to calculate the 3D reweighted PMF and associated ligand binding free energies.

### Ligand binding kinetics obtained from reweighting of LiGaMD2 Simulations

Reweighting of ligand binding kinetics from LiGaMD2 simulations followed a similar protocol using Kramers’ rate theory that has been recently implemented in kinetics reweighting of the GaMD^35^, LiGaMD^34b^, Pep-GaMD^29^ and PPI-GaMD^36^ simulations. Provided sufficient sampling of repetitive ligand dissociation and binding in the simulations, we record the time periods and calculate their averages for the ligand found in the bound (*τ*_*B*_) and unbound (*τ*_*U*_) states from the simulation trajectories. The *τ*_*B*_ corresponds to residence time in drug design^37^. Then the ligand dissociation and binding rate constants (*k*_*off*_ and *k*_*on*_) were calculated as:

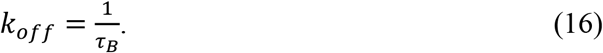

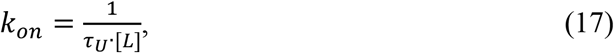

where [L] is the ligand concentration in the simulation system.

According to Kramers’ rate theory, the rate of a chemical reaction in the large viscosity limit is calculated as^35^:

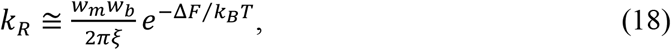

where *w*_*m*_ and *w*_*P*_ are frequencies of the approximated harmonic oscillators (also referred to as curvatures of free energy surface^38^) near the energy minimum and barrier, respectively, *ξ* is the frictional rate constant and Δ*FF* is the free energy barrier of transition. The friction constant *ξ* is related to the diffusion coefficient *D* with *ξ*= *k*_*B*_*T*/*D*. The apparent diffusion coefficient *D* can be obtained by dividing the kinetic rate calculated directly using the transition time series collected directly from simulations by that using the probability density solution of the Smoluchowski equation^39^. In order to reweight ligand kinetics from the LiGaMD2 simulations using the Kramers’ rate theory, the free energy barriers of ligand binding and dissociation are calculated from the original (reweighted, **Δ*F***) and modified (no reweighting, **Δ*F\****) PMF profiles, similarly for curvatures of the reweighed (*w*) and modified (*w*^*^, no reweighting) PMF profiles near the ligand bound (“B”) and unbound (“U”) low-energy wells and the energy barrier (“Br”), and the ratio of apparent diffusion coefficients from simulations without reweighting (modified, *D*^*^) and with reweighting (*D*). The resulting numbers are then plugged into Eq. (17) to estimate accelerations of the ligand binding and dissociation rates during LiGaMD2 simulations^35^, which allows us to recover the original kinetic rate constants.

### System Setup

The complex structures of benzene (BEN) bound to the L99A T4L, M102A T4L and F104A T4L were obtained from the 3HH4^40^, 220L^41^ and 227L^41^ PDB files, respectively. The crystal structure of indole (IND) bound to the L99A T4L was obtained from the 185L^42^ PDB file. The AMBER ff14SB force field^43^ was used for the protein and the GAFF force field^44^ for the ligand with AM1-BCC charges. Each system was solvated in a periodic box of TIP3P water molecules with a distance of 18 Å from the solute to the box edge using *tleap*. Therefore, the ligand concentration was 0.0034 M in the simulation system. Appropriate number of Clions were added to achieve system neutrality.

### Simulation Protocol

Each system was energy minimized and gradually heated to 300 K in 1 ns with the Langevin thermostat and harmonic restraints of 50 kcal/mol/Å^2^ on all non-hydrogen atoms of the protein and the ligand using the AMBER22 software^45^. The simulation system was firstly energy minimized with 1.0 kcal/mol/Å^2^ constraints on the heavy atoms of the proteins, including the steepest descent minimization for 50,000 steps and conjugate gradient minimization for 50,000 steps. The system was then heated from 0 K to 300 K for 200 ps. It was further equilibrated using the NVT ensemble at 300 for 200 ps and the NPT ensemble at 300 K and 1 bar for 1 ns with 1 kcal/mol/Å^2^ constraints on the heavy atoms of the protein, followed by 2 ns short cMD without any constraint. The LiGaMD2 simulations proceeded with 14 ns short cMD to collect the potential statistics, 49.2 ns LiGaMD2 equilibration after adding the boost potential and then three independent 1,000 ns production runs. It provided more powerful sampling to set the threshold energy for applying the boost potential to the upper bound (i.e., *E* = *V*_min_+1/*k*) in our previous study ligand dissociation and binding using LiGaMD^29^. Therefore, the threshold energy for applying the ligand essential potential was also set to the upper bound in the LiGaMD2 simulations. The selective boost potential was applied to both the ligand and protein pocket residues within 5Å of ligand in the LiGaMD2 simulations. For the second boost potential applied on the system total potential energy other than the ligand essential potential energy, sufficient acceleration was obtained by setting the threshold energy to the lower bound. In order to observe ligand dissociation during LiGaMD2 equilibration while keeping the boost potential as low as possible for accurate energetic reweighting, the (σ_0*P*_, σ_0*D*_) parameters were finally set to (9.0 kcal/mol, 6.0 kcal/mol), (7.0 kcal/mol, 6.0 kcal/mol), (9.0 kcal/mol, 6.0 kcal/mol) and (9.0 kcal/mol, 6.0 kcal/mol) for the LiGaMD2 simulations of the BEN bound to the L99A T4L (T4L:L99A-BEN), F104A T4L (T4L:F104A-BEN), M102A T4L (T4L:M102A-BEN) and IND bound to the L99A T4L (T4L:L99A-IND). LiGaMD2 production simulation frames were saved every 0.4 ps for analysis.

### Simulation Analysis

The VMD^46^ and CPPTRAJ^47^ tools were used for simulation analysis. The number of ligand dissociation and binding events (*N*_*D*_ and *N*_*B*_) and the ligand binding and unbinding time periods (*τ*_*B*_ and *τ*_*U*_) were recorded from individual simulations (**Tables 1 &S1**). With high fluctuations, *τ*_*B*_ and *τ*_*U*_ were recorded for only the time periods longer than 1 ns. The 1D, 2D and 3D PMF profiles, as well as the ligand binding free energy, were calculated through energetic reweighting of the LiGaMD2 simulations. The center-of-mass distance between the ligand and the protein pocket (defined by protein residues within 5 Å of ligand) and ligand heavy atom RMSDs relative to X-ray structures with the protein aligned were chosen as the reaction coordinate for calculating the 1D PMF profiles. The bin size was set to 1.0 Å. The software *trj_cavity*^*48*^ implemented in GROMACS was used to calculate the pocket volume. 2D PMF profiles of the ligand RMSD and pocket volume were calculated to analyze conformational changes of the protein upon ligand binding. The bin size was set to 50 Å^3^ for pocket volume. The cutoff for the number of simulation frames in one bin was set to 500 for reweighting 1D and 2D PMF profiles. The 3D PMF profiles of ligand displacements from the T4L mutants in the X, Y and Z directions were further calculated from the LiGaMD2 simulations. The bin sizes were set to 1 Å in the X, Y and Z directions. The cutoff of simulation frames in one bin for 3D PMF reweighting (ranging from 100-400 for three individual LiGaMD2 simulations) was set to the minimum number below which the calculated 3D PMF minimum will be shifted. The ligand binding free energies (Δ*G*) were calculated using the reweighted 3D PMF profiles and binding kinetic rates by Δ*G* = −RTLn*k*_*off*_/*k*_*on*_, respectively. The ligand dissociation and binding rate constants (*k*_*on*_ and *k*_*off*_) were calculated from the LiGaMD2 simulations with their accelerations analyzed using the Kramers’ rate theory (**Table S2**).

**Table 1.** Summary of LiGaMD2 simulations performed on ligand binding to the T4L mutants. Δ*V* is the total boost potential. *N*_*D*_ and *N*_*B*_ are the number of observed ligand dissociation and binding events, respectively. Δ*G*_*sim*_ and Δ*G*_*exp*_ are the ligand-T4L binding free energies obtained from LiGaMD2 simulations and experiments, respectively. 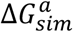 and 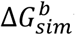 were calculated with 3D PMF and reweighted binding kinetic rates by Δ*G* = −RTLn (*k*_*off*_/*k*_*on*_, respectively.

| T4L | Ligand | ID | N<sub>B</sub> | N<sub>D</sub> | $\Delta V$ (kcal/mol) | $\Delta G_{sim}^a$<br>(kcal/mol) | $\Delta G_{sim}^b$<br>(kcal/mol) | $\Delta G_{exp}$<br>(kcal/mol) |
| --- | --- | --- | --- | --- | --- | --- | --- | --- |
| L99A | BEN | Sim1 | 6 | 6 | 107.77±9.67 | -5.88±0.61 | -5.17±0.72 | -4.12 |
|  |  | Sim2 | 6 | 6 | 109.84±9.58 |  |  |  |
|  |  | Sim3 | 5 | 5 | 109.63±9.59 |  |  |  |
| M102A | BEN | Sim1 | 6 | 6 | 108.10±10.04 | -6.43±0.12 | -5.01±0.73 | -4.18 |
|  |  | Sim2 | 3 | 3 | 108.04±9.97 |  |  |  |
|  |  | Sim3 | 7 | 7 | 109.21±9.49 |  |  |  |
| F104A | BEN | Sim1 | 4 | 4 | 70.96±8.11 | -6.02±0.70 | -3.42±0.72 | -4.02 |
|  |  | Sim2 | 3 | 3 | 72.53±7.84 |  |  |  |
|  |  | Sim3 | 3 | 3 | 70.45±7.81 |  |  |  |
| L99A | IND | Sim1 | 6 | 7 | 115.80±9.68 | -7.40±0.39 | -4.87±1.06 | -4.82 |
|  |  | Sim2 | 3 | 3 | 118.16±9.79 |  |  |  |
|  |  | Sim3 | 4 | 4 | 116.49±9.71 |  |  |  |

## Results

### Flexibity of the protein pocket plays an important role in dissociation of buried ligands

The ligand binding pockets are different in the L99A and F104A mutants of T4 lysozyme (T4L). The binding pocket in L99A T4L is deeply buried (**Fig. 1A**), while the pocket is exposed to the solvent in the F104A mutant (**Fig. 1D**). LiGaMD and LiGaMD2 testing simulations with *σ*_0*P*_increasing from 6.0 to 10.0 kcal/mol and *σ*_0*D*_ at 6.0 kca/mol were performed on these two systems. Ligand dissociation was captured in both LiGaMD and LiGaMD2 simulations of benzene (BEN) binding in the F104A mutant with an exposed binding pocket (**Fig. 1E& 1F**). With LiGaMD, the ligand dissociated from the F104A T4L at 5.45 ns, 7.20 ns, 18.22 ns and 7.80 ns with the *σ*_0*D*_ values of 6.0, 7.0, 8.0 and 9.0 kcal/mol, respectively. Interestingly, LiGaMD with the *σ*_0*P*_ value at 9.0 could capture both the ligand dissociation and rebinding within the 49.2 ns equilibration simulation. LiGaMD2 captured the ligand dissociation at 3.60 ns, 11.20 ns, 4.50 ns and 29.60 ns with the *σ*_0*P*_ value of 6.0, 7.0, 8.0 and 9.0 kcal/mol, respectively. The LiGaMD2 with the *σ*_0*P*_ values of 7.0 and 8.0 kcal/mol could capture the ligand dissociation and rebinding within the 49.2 ns equilibration simulation.

**Figure 1.**
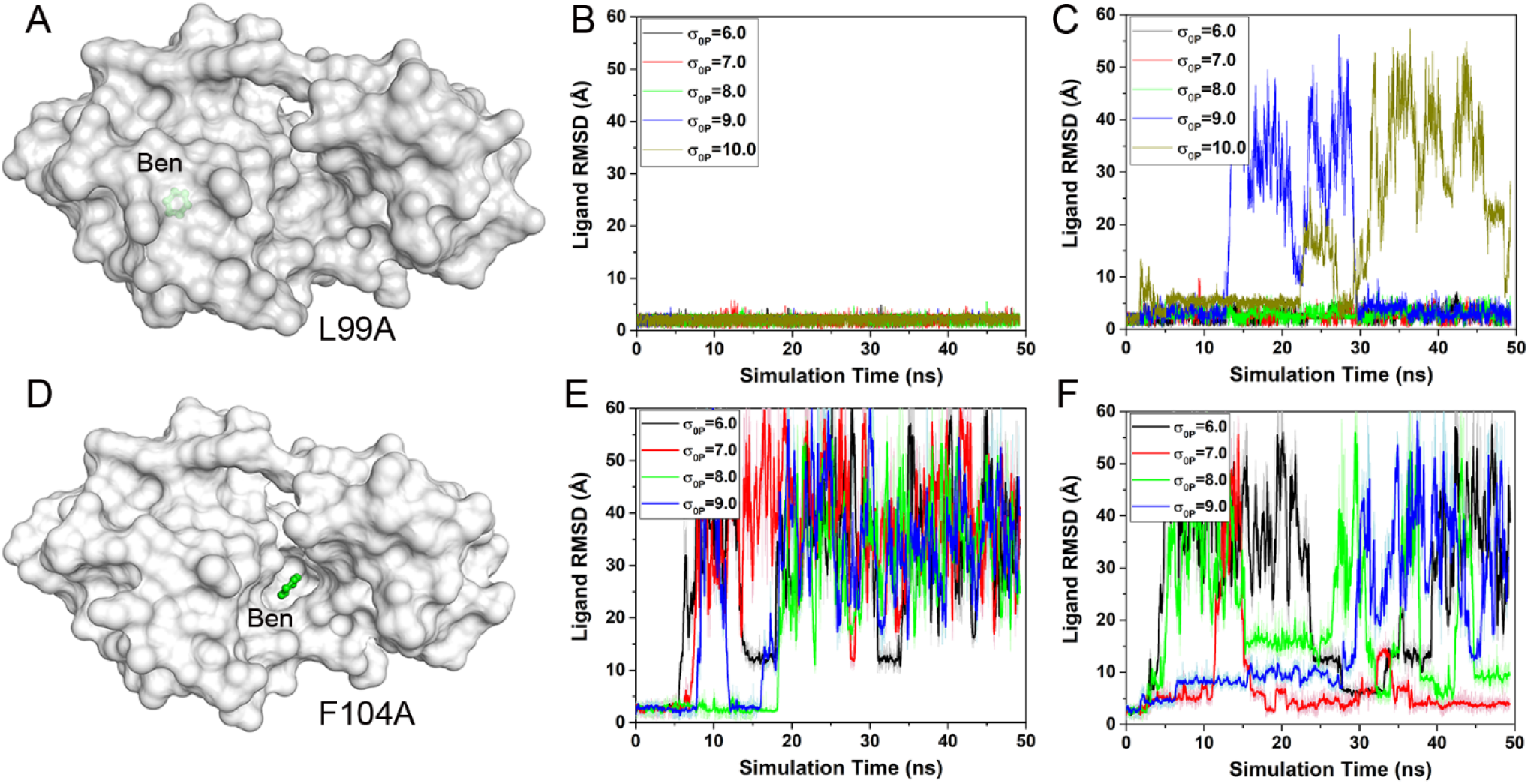
Comparison of LiGaMD and LiGaMD2 simulations on the T4L mutant systems with buried and open binding pockets: Computational models of benzene binding to the T4L:L99A with a burred binding pocket (**A**) and T4L:F104A with an open binding pocket (**D**); Time courses of ligand root-mean-square deviation (RMSD) in T4L:L99A calculated from 49.2 ns LiGaMD (**B**) and LiGaMD2 (**C**) equilibration simulations, respectively. Time courses of ligand RMSD in F104A T4L calculated from 49.2 ns LiGaMD (**E**) and LiGaMD2 (**F**) equilibration simulations, respectively.

For the L99A T4L with a buried protein pocket, the LiGaMD could not capture ligand dissociation even with the value of *σ*_0*P*_ increased to 10.0 kcal/mol (**Fig. 1B**). In comparison, the LiGaMD2 simulations could capture the ligand dissociation and rebinding with the *σ*_0*P*_ values of 9.0 and 10.0 kcal/mol (**Fig. 1B&1C**). The ligand dissociated at 12.52 ns and rebound at ∼29.50 ns in the LiGaMD2 simulation with the *σ*_0*P*_ value of 9.0 kcal/mol. In the LiGaMD2 simulation with the *σ*_0*P*_ value of 10.0 kcal/mol, the ligand dissociation and rebinding occurred at ∼23.60 ns and ∼26.80 ns, respectively.

Next, we identified correlation of the ligand dissociation and conformational changes of the protein pocket in the L99A T4L (**Fig. S1**). The binding pocket exhibited low RMSD of ∼1-2 Å in the LiGaMD simulations with all the tested parameters as no selective boost potential was applied to the protein pocket (**Fig. S1A**). In comparison, RMSD of the binding pocket significantly increased in the LiGaMD2 simulations with the *σ*_0*P*_ values of 9.0 and 10.0 kcal/mol, respectively (**Fig. S1B**). As the *σ*_0*P*_ value of 9.0 kcal/mol was the lowest value that enabled the ligand dissociation and rebinding, we further focused on the binding pocket in this simulation. During ligand dissociation around 12 ns, RMSD of the binding pocket increased to ∼1.7 Å. RMSD of the binding pocket decreased to mostly <1.0 Å when the ligand completely dissociated to the solvent. The binding pocket RMSD increased to ∼2 Å when the ligand rebound to the pocket at ∼30 ns (**Fig S1C**). After the ligand bound completely to the protein pocket, RMSD of the binding pocket deceased to ∼1 Å again. We further calculated volume of the protein pocket during the LiGaMD2 simulation (**Fig S1D**). The volume of the protein pocket was overall smaller in the ligand-free (*apo*) state than in the ligand-bound (*holo*) state (**Fig S1D**). Similar opening of the binding pocket was observed in previous aMD simulation of BEN dissociation from the L99A T4L^49^. In summary, LiGaMD2 showed improved sampling of ligand binding to the buried protein pockets, where dynamics of the binding pocket played an important role in the ligand dissociation and rebinding. For the system with an open pocket, both LiGaMD and LiGaMD2 worked well.

### Microsecond LiGaMD2 simulations captured repetitive ligand dissociation and rebinding to the T4L mutants

As the *σ*_0*P*_values of 9.0 and 7.0 kcal/mol were the lowest to capture the ligand dissociation and rebinding in LiGaMD2 equilibration simulations of the L99A and F104A T4L systems (**Figs. 1C & 1F**), they were used for further three independent 1,000 ns production simulations. Furthermore, two more systems with buried binding pockets were added to demonstrate the robustness of LiGaMD2, including the M102A T4L bound by BEN and the L99A T4L bound by a different ligand indole (IND). In summary, LiGaMD2 simulations were performed on three complexes of BEN bound to the L99A T4L (T4L:L99A-BEN), M102A T4L (T4L:M102A-BEN) and F104A T4L (T4L:F104A-BEN), and another complex of IND bound to the L99A T4L (T4L:L99A-IND) (**Figs. 2A-D**). Three independent 1,000 ns LiGaMD2 production simulations were performed on each of the four systems (**Table 1**). The LiGaMD2 simulations of the T4L:L99A-BEN system recorded an average boost potential of 107.77-109.84 kcal/mol with 9.58-9.67 kcal/mol standard deviation. The LiGaMD2 simulations of the T4L:M102A-BEN system recorded an average boost potential of 108.04-109.21 kcal/mol with 9.49-10.04 kcal/mol standard deviation. In comparison, the average boost potential was 70.45-72.53 kcal/mol with 7.81-8.11 kcal/mol standard deviation in the three simulations of the T4L:F104A-BEN. The boost potential applied in simulations of the T4L:L99A-IND system was slightly larger than that of the T4L:L99A-BEN system, with average of 115.80-118.16 kcal/mol and 9.68-9.79 kcal/mol standard deviation (**Table 1**).

**Figure 2.**
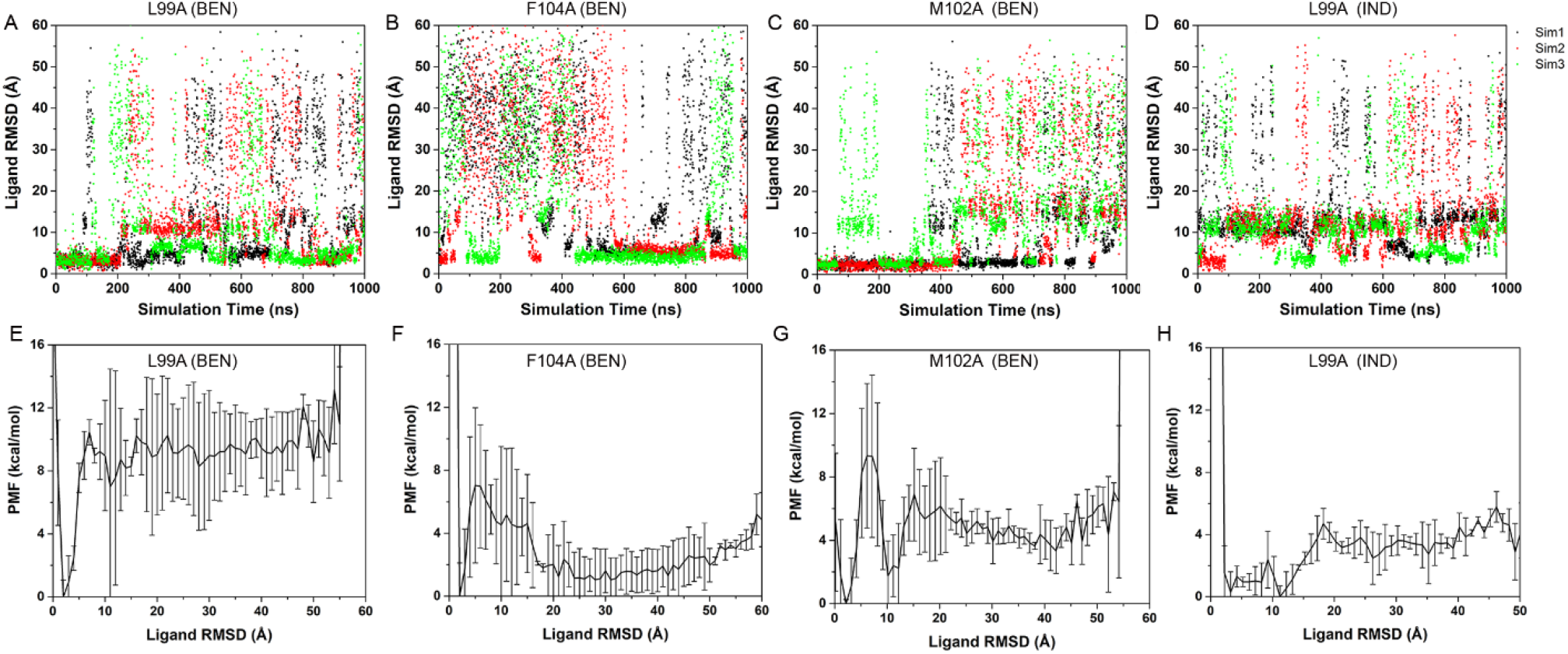
LiGaMD2 simulations captured repetitive dissociation and binding of two different ligands (benzene and indole) to T4L mutants: (**A**-**D**) time courses of ligand heavy atom RMSDs relative to X-ray structures calculated from three independent 1 μs LiGaMD2 simulations of (**A**) benzene binding to T4L:L99A, (**B**) benzene binding to T4L:F104A, (**C**) benzene binding to the T4L:M102A and (**D**) indole binding to the T4L:L99A. (**E**-**H**) The corresponding PMF profiles of the ligand RMSDs averaged over three LiGaMD2 simulations of (**E**) benzene binding to T4L:L99A, (**F**) benzene binding to T4L:F104A, (**G**) benzene binding to the T4L:M102A and (**H**) indole binding to the T4L:L99A. Error bars are standard deviations of the free energy values calculated from three LiGaMD2 simulations.

RMSDs of the ligand relative to the X-ray structures with the T4L aligned were calculated (**Figs. 2A-2D**) to record the number of ligand dissociation and binding events (*N*_*D*_ and *N*_*B*_) in each of the 1,000 ns LiGaMD2 simulations. With close examination of the ligand binding trajectories, RMSD cutoffs of the ligand unbound and bound states were set to >15 Å and <5.0 Å, respectively. Because of ligand fluctuations, we recorded only the corresponding binding and dissociation events that lasted for more than 1.0 ns. In each simulation of the T4L:L99A-BEN system, about 5-6 binding and 5-6 dissociation events were observed (**Fig. 2A & Table 1**). In each simulation of the T4L:F104A-BEN system, about 3-4 dissociation and 3-4 binding events were observed (**Fig. 2B & Table 1**). In each simulation of the T4L:M102A-BEN system, about 3-7 dissociation and 3-7 binding events were observed (**Fig. 2C & Table 1**). A similar number of ligand dissociation (3-6) and binding (3-7) events were observed in simulations of the T4L:L99A-IND system (**Fig. 2D & Table 1**). In summary, repetitive ligand dissociation and rebinding were successfully captured in each of the 1,000 ns LiGaMD2 simulations of various T4L mutants bound by different ligands (**Figs. 2**).

Next, we explored the correlation between conformational changes of the binding pocket and ligand binding in the LiGaMD2 production simulations. The ligand RMSD and volume of the binding pocket were used as reaction coordinates to calculate 2D PMF (**Fig. 3**). Five low-energy states were identified in the 2D PMF profile of the T4L:L99A-BEN system including the Bound (“B”), Intermediate (“I”), Unbound U1 (“U1”), Unbound U2 (“U2”) and Unbound U3 (“U3”) states (**Fig. 3A**). The ligand RMSD and pocket volume in the B, I, U1, U2 and U3 states centered around (2.5 Å, 300 Å^3^), (9.3 Å, 425.7 Å^3^), (24.5 Å, 259.9 Å^3^), (34.8 Å, 360.3 Å^3^), and (41.6 Å, 251.9 Å^3^), respectively (**Fig 3A**). In the T4L:F104A-BEN system, three low-energy states were identified, including the Bound (“B”), Intermediate (“I”) and Unbound (“U”) states (**Fig. 3B**). The ligand RMSD and the pocket volume in the B, I and U centered around (3.8 Å, 176.80 Å^3^), (11.6 Å, 59.6 Å^3^) and (35.0 Å, 54.7 Å^3^), respectively (**Fig. 3B**). In the T4L:M102A-BEN system, five low-energy states were identified including the Bound (“B”), Intermediate (“I”), Intermediate (“I2”), Unbound (“U1”) and Unbound (“U2”) states (**Fig. 3C**). The ligand RMSD and pocket volume in the B, I, I2, U1 and U2 states centered around (2.7 Å, 219.7 Å^3^), (10.7 Å, 356.5 Å^3^), (11.0 Å, 113.4 Å^3^), (32.9 Å, 258.8 Å^3^), and (37.8 Å, 132.9 Å^3^), respectively (**Fig 3C**). In the T4L:L99A-IND system, four low-energy states were identified including the Bound (“B”), Intermediate (“I”), Intermediate (“I2”) and Unbound (“U”) states (**Fig. 3D**). The ligand RMSD and pocket volume in the B, I, I2, and U states centered around (1.9 Å, 363.2 Å^3^), (5.0 Å, 468.8 Å^3^), (11.5 Å, 357.4 Å^3^), and (28.8 Å, 155.4 Å^3^), respectively (**Fig 3D**). In compared with the bound state, a larger binding pocket volume was identified in the intermediate I state in the systems with an burred protein pocket including the T4L:L99A-BEN, T4L:M102A-BEN and T4L:L99A-IND (**Figs. 3A, 3B & 3D**). In contrast, the binding pocket volume was significantly smaller in the intermediate I state in the T4L:F104A-BEN with an exposed binding pocket (**Fig. 3B**). The representative intermediate conformational states from the four systems were shown in **Fig. 3E-3H**. Compared to the bound X-ray structures, helices C and G in the intermediate “I” state moved outward in the T4L:L99A-BEN system, leading to opening of the binding pocket (**Fig. 3E**). In the T4L:F104A-BEN system, major conformational changes upon ligand binding involved helices C and B, which moved inwards and reduced volume of the binding pocket (**Fig. 3F**). In the T4L:M102A-BEN system, helices D, F and G moved outwards in the intermediate “I” state with opening of the ligand binding pocket, accompanied by inward movement of helix C (**Fig. 3G**). In the T4L:L99A-IND system, outward movements were observed in helices C, D and G in the intermediate I state (**Fig. 3H**). Therefore, conformational changes of the protein pocket greatly facilitated the ligand dissociation and binding in the L99A and M104A systems with a buried binding pocket.

**Figure 3.**
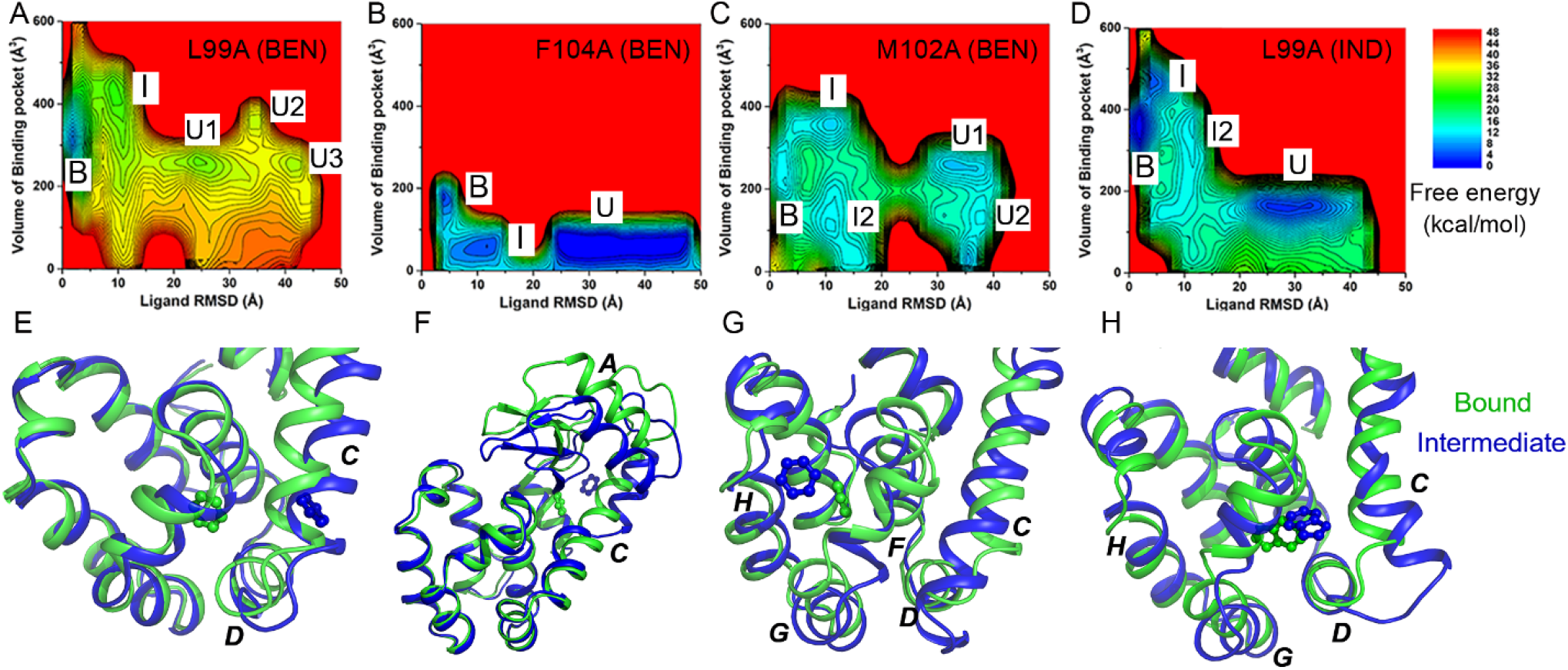
2D Free energy profiles and low-energy intermediate conformational states of ligand binding to the T4L mutants: (**A-D**) 2D PMF profiles regarding the ligand heavy atom RMSD and the pocket volume in LiGaMD2 simulations of (**A**) benzene binding to T4L:L99A, (**B**) benzene binding to T4L:F104A, (**C**) benzene binding to T4L:M102A, (**D**) indole binding to T4L:F99A. (**E**-**H**) Low-energy “Intermediate” (“I”) conformations (blue) as identified from the 2D PMF profiles of (**E**) benzene binding to T4L:L99A, (**F**) benzene binding to T4L:F104A, (**G**) benzene binding to T4L:M102A and (**H**) indole binding to T4L:L99A. X-ray structures of the ligand-bound complexes (“Bound”) are shown in green. The ligands are shown in balls and sticks, and the helix are shown in cartoon. The helix C, D, F and G are labeled as they show significant changes between the “Bound” and “Intermediate” conformational states.

### Ligand binding kinetic rates and free energies calculated from LiGaMD2 agreed well with experimental data

LiGaMD2 simulations that successfully captured repetitive ligand binding and dissociation allowed us to calculate the ligand binding kinetic rate constants. We recorded the time periods for the ligand found in the bound (*τ*_*B*_) and unbound (*τ*_*U*_) states throughout the LiGaMD2 simulations. Without reweighting, the ligand binding rate constants (*k*_*on*_*\**) were calculated directly from the LiGaMD2 trajectories as 8.22 ± 5.48×10^7^ M^-1^·s^-1^, 7.79±1.36 × 10^6^ M^-1^·s^-1^, 6.81±1.27×10^7^ M^-1^·s^-1^ and 1.67±0.67×10^9^ M^-1^·s^-1^ in the system of T4L:L99A-BEN, T4L:M102A-BEN, T4L:F104A-BEN and T4L:L99A-IND systems, respectively (**Table 2**). The accelerated dissociation rate constants (*k*_*off*_*\**) were calculated as 3.47 ± 2.31×10^5^ s^-1^, 3.46 ±1.81×10^9^ s^-1^, 1.72±1.44×10^9^ s^-1^ and 2.49±1.25×10^7^ s^-1^ in the T4L:L99A-BEN, T4L:M102A-BEN, T4L:F104A-BEN and T4L:L99A-IND systems, respectively (**Table 2**).

**Table 2.** Comparison of kinetic rates obtained from experiments and LiGaMD2 simulations for ligand binding to T4L mutants. *k*_*on*_ and *k*_*off*_ are the kinetic dissociation and binding rate constants, respectively, from experimental data or LiGaMD2 simulations with reweighting using Kramers’ rate theory. *k*_*on*_*\** and *k*_*off*_*\** are the accelerated kinetic dissociation and binding rate constants calculated directly from LiGaMD2 simulations without reweighting.

| System | Method | $k_{on} (M^{-1} \cdot s^{-1})$ | $k_{off} (s^{-1})$ | $k_{on}^* (M^{-1} \cdot s^{-1})$ | $k_{off}^* (s^{-1})$ |
| --- | --- | --- | --- | --- | --- |
| T4L:L99A-BEN | Experiment | $0.8-1.0 \times 10^6$ | $9.50 \times 10^2$ | - | - |
| | LiGaMD2 | $7.42 \pm 4.81 \times 10^6$ | $1.44 \pm 0.88 \times 10^3$ | $8.22 \pm 5.48 \times 10^7$ | $3.47 \pm 2.31 \times 10^5$ |
| T4L:M102A-BEN | Experiment | $3.0-5.0 \times 10^6$ | $3.00 \times 10^3$ | | |
| | LiGaMD2 | $9.57 \pm 6.29 \times 10^6$ | $2.01 \pm 1.61 \times 10^3$ | $7.79 \pm 1.36 \times 10^6$ | $3.46 \pm 1.81 \times 10^9$ |
| T4L:F104A-BEN | Experiment | $>10 \times 10^6$ | $>1.00 \times 10^4$ | | |
| | LiGaMD2 | $3.16 \pm 2.29 \times 10^9$ | $1.38 \pm 0.67 \times 10^6$ | $6.81 \pm 1.27 \times 10^7$ | $1.72 \pm 1.44 \times 10^9$ |
| T4L:L99A-IND | Experiment | $0.7-1.0 \times 10^6$ | $3.25 \times 10^2$ | | |
| | LiGaMD2 | $2.99 \pm 2.87 \times 10^6$ | $3.49 \pm 0.56 \times 10^3$ | $1.67 \pm 0.67 \times 10^9$ | $2.49 \pm 1.25 \times 10^7$ |

Next, we reweighted the LiGaMD2 simulations of ligand-T4L mutants to calculate acceleration factors of ligand binding and dissociation processes (**Table S1**) and recovered the original kinetic rate constants using the Kramers’ rate theory (**Table 2**). The dissociation free energy barrier (Δ*F*_*off*_) significantly decreased from 10.5±0.86, 9.00±1.07, 8.55±0.54, and 7.11±0.34 kcal/mol in the reweighted PMF profiles to 2.55±0.43, 2.35±0.34, 2.36±0.35 and 2.23±0.26 kcal/mol in the modified PMF profiles for the system of T4L:L99A-BEN, T4L:M102A-BEN, T4L:F104A-BEN and T4L:L99A-IND, respectively, respectively (**Table S1 and Fig. S2**). The free energy barrier for ligand binding (Δ*F*_*on*_) slightly decreased from 4.93±1.38, 6.03±0.11, 5.40±1.07, 4.37±0.33 kcal/mol in the reweighted profiles to 0.57±0.14, 1.15±0.50, 0.54±0.053, 0.53±0.10 kcal/mol in the modified PMF profiles for the system of T4L:L99A-BEN, T4L:M102A-BEN, T4L:F104A-BEN and T4L:L99A-IND, respectively (**Table S1 and Fig. S2**). Curvatures of the reweighed (*w*) and modified (*w*^*^, no reweighting) free energy profiles were calculated near the ligand Bound (“B”) and Unbound (“U”) low-energy wells and the energy barrier (“Br”), as well as the ratio of apparent diffusion coefficients calculated from LiGaMD2 simulations with reweighting (*D*) and without reweighting (modified, *D*^*^) (**Table S1**). According to the Kramers’ rate theory, the ligand association was accelerated by 11.07, 0.81, 0.02 and 558.5 times for the systems of T4L:L99A-BEN, T4L:M102A-BEN, T4L:F104A-BEN and T4L:L99A-IND, respectively. The ligand dissociation was significantly accelerated by 2.40×10^2^, 1.72×10^6^, 1.24×10^3^ and 7.13×10^3^ times in the T4L:L99A-BEN, T4L:M102A-BEN, T4L:F104A-BEN and T4L:L99A-IND systems, respectively. Therefore, the reweighted *k*_*on*_ in the T4L:L99A-BEN, T4L:M102A-BEN, T4L:F104A-BEN and T4L:L99A-IND systems were calculated as 7.42±4.81×10^6^, 9.57±6.29×10^6^, 3.16±2.29×10^9^ and 2.99±2.87×10^6^ M^-1^·s^-1^, being highly consistent with the corresponding experimental values^50^ of 0.8-1.0×10^6^, 3.0-5.0×10^6^, >10×10^6^ and 0.7-1.0×10^6^ M^-1^·s^-1^, respectively (**Table 2**). The reweighted *k*_*off*_ in the T4L:L99A-BEN, T4L:M102A-BEN, T4L:F104A-BEN and T4L:L99A-IND systems were calculated as 1441±883, 2011±1606, 1.38±0.67×10^6^ and 3494±559 s^-1^, in good agreement with the corresponding experimental values^50^ of 950, 3000, >10000 and 325 s^-1^, respectively.

Based on the ligand binding kinetic rates (*k*_*on*_ and *k*_*off*_), we calculated the ligand binding free energies as Δ*G* = −RTLn*k*_*off*_/*k*_*on*_. The resulting binding free energies in the T4L:L99A-BEN, T4L:M102A-BEN, T4L:F104A-BEN and T4L:L99A-IND systems (**Table 1**) were -5.17±0.72 kcal/mol, -5.01±0.73 kcal/mol, -3.42±0.72 kcal/mol and -4.87±1.06 kcal/mol, being highly consistent with the corresponding experimental values of -4.12 kcal/mol, -4.18 kcal/mol, -4.02 kcal/mol and -4.82 kcal/mol, respectively. The root-mean square error (RMSE) of binding free energy was only 0.73 kcal/mol. Alternatively, we could calculate the ligand binding free energy using the 3D reweighted PMF profiles of the ligand displacement from the T4L binding pocket in the X, Y and Z directions (**Table 1**). The ligand binding free energies in the T4L:L99A-BEN, T4L:M102A-BEN, T4L:F104A-BEN and T4L:L99A-IND systems (**Table 1**) were estimated as - 5.88±0.61 kcal/mol, -6.43±0.70 kcal/mol, -4.57± 0.12 kcal/mol and -7.40±0.39 kcal/mol, respectively. The RMSE of binding free energies predicted from the 3D PMF was greater as 1.94 kcal/mol, but still within the acceptable range of binding free energy predictions (2.0 kcal/mol)^51^. Therefore, both efficient conformational sampling and accurate ligand binding thermodynamic and kinetic calculations were achieved through the LiGaMD2 simulations.

### Multiple ligand binding and dissociation pathways were identified from LiGaMD2 simulations

We closely examined the LiGaMD2 trajectories to explore ligand binding and dissociation pathways. For dissociation of BEN from the T4L:L99A, four pathways between the CD, CF, DG and FGH helices were identified (**Fig. 4**). All these pathways were involved in the BEN rebinding to the T4L:L99A. One extra binding pathway of HG was found in the BEN binding to the T4L:L99A. LiGaMD2 captured 5, 3, 6 and 3 times of BEN dissociation events through pathways of CD, CF, DG and FGH, respectively (**Fig 4**). The BEN rebinding events through pathways of CD, CF, DG, FGH and HG were 3, 2, 3, 6 and 2, respectively. The same ligand dissociation pathways of CD, CF, DG and FGH were identified in the LiGaMD2 simulations of T4L:M102A-BEN and T4L:L99A-IND systems (**Fig. 4**). Two pathways of DG and FGH were observed in the simulations of BEN binding to the T4L:M102A. The ligand dissociation events in T4L:M102A system along pathways of CD, CF, DG and FGH were 1, 2, 1 and 12, respectively (**Fig 4**). The ligand rebinding events in the T4L:M102A-BEN through pathways of DG and FGH were 2 and 11, respectively. For dissociation of BEN from the T4L:F104A, two pathways near the A and C helices were identified. The dissociation events in the T4L:F104A-BEN were 8 and 1 via pathways A and C, respectively (**Fig. 4**). Pathway C was observed in BEN binding to the F104A T4L mutant (**Fig. 4**). For the dissociation of IND from the T4L:L99A, four pathways between the CD, CF, DG and FGH helices were identified. While only pathways CF and FGH were observed in the rebinding of IND to the L99A T4L mutant. The dissociating event via pathways CD, CF, DG and FGH in the T4L:L99A-IND system were 3, 3, 5, and 3, respectively. The IND binding events via pathways CF and FGH were 3 and 9, respectively (**Fig. 4**). The binding and dissociating pathways were consistent with earlier simulation findings using Metadynamics^28, 52^, Weighted Ensemble^20b^, Machine Learning^53^, tRAMD^15a^ and aMD^49^ simulations. In summary, multiple ligand binding and dissociation pathways were observed in the LiGaMD2 simulations of the T4L mutants. The ligand binding and dissociation mostly followed the same pathways.

**Figure 4.**
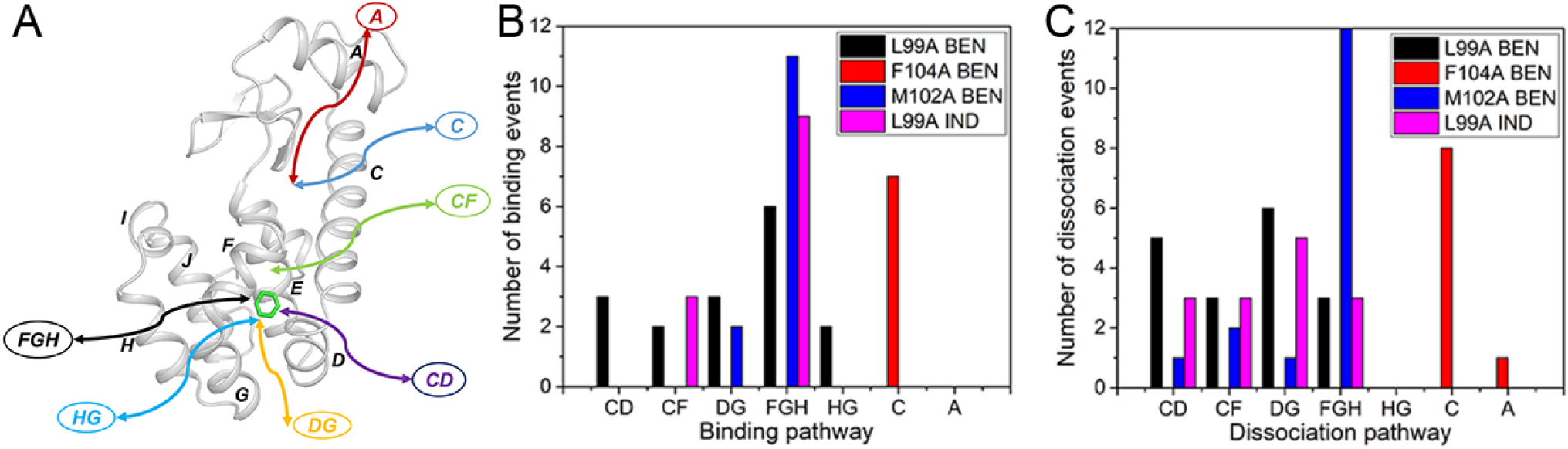
Pathways of ligand binding and dissociation in the T4L mutants. (**A**) Cartoon representation of the protein with helices labelled. Binding and dissociation pathways are denoted by the arrow lines. Number of binding (**B**) and dissociation (**C**) events through the different pathways captured by the LiGaMD2 simulations.

## Discussions

We have presented LiGaMD2 that improved enhanced sampling and accurate prediction of protein-ligand binding thermodynamics and kinetics for especially proteins with closed binding pockets. LiGaMD2 works by selectively boosting the potential of both ligand and protein residues in the binding pocket. LiGaMD2 shows significantly improved sampling of systems with buried binding pockets, where flexibility of the binding pocket plays a critical role in ligand binding. Microsecond LiGaMD2 simulations have allowed us to capture repetitive ligand dissociation and rebinding processes as demonstrated on four T4L mutant model systems. These simulations then enabled accurate predictions of ligand binding free energies and kinetic rate constants.

LiGaMD2 simulations revealed the critical role of protein flexibility for ligand binding, especially in the case of solvent-inaccessible buried pockets, in good agreement with previous experimental^52a, 54^ and computational studies^14d, 20b, 49^. Protein flexibility has been recognized as one of the main factors that regulates protein-ligand binding kinetics^25c-e^. The influence of protein flexibility on ligand binding site can vary from small changes like opening or closing of an existing pocket to the formation of a new pocket^25e^. For example, the MSM^8^ built with 60 μs cMD simulations revealed that the movement of helix D/G/H/J could transiently open a channel for ligand binding to the target site of the L99A T4L. Such movement of helix D/G/H were also observed in the intermediate states in the LiGaMD2 simulations (**Fig. 3**). Additionally, multiple ligand binding and dissociation pathways were identified from LiGaMD2 simulations (**Fig. 4**), being highly consistent with previous enhanced sampling simulations, including the RAMD^15a^, aMD^49^, Metadynamics^28^, MSM^8^ and Weighted Ensemble^20b^. For example, the dissociation pathway FGH with highest probability observed in LiGaMD2 was also captured in the simulations of RAMD^15a^, Metadynamics^28^, aMD^49^, MSM^8^ and Weighted Ensemble^20b^.

Compared with the cMD^55^, Metadynamics^56^, Weighted Ensemble,^57^ MSM^8^ and Replica Exchange MD simulations^25a^, LiGaMD2 provides an efficient and/or easier-to-use approach to simulation of ligand binding and dissociation and calculations of ligand binding thermodynamics and kinetics. It is advantageous over previous LiGaMD for proteins with buried binding pockets. Microsecond cMD simulations were able to capture benzene binding to the L99A T4L^8^. However, slower ligand dissociation was still beyond the accessibility of cMD. Weighted Ensemble^20b^ and MSM were able to accurately predict ligand binding kinetics^8^. However, tens of microsecond cMD simulations were needed for the Weighted Ensemble^20b^ and MSM^8^. For the Replica Exchange method^58^, a large number of replica simulations were often needed to model protein-ligand binding. In the case of gREST simulations^14d^, eight replicas were needed to capture ligand binding to the L99A T4L. With carefully designed CV, Metadynamics could capture both ligand binding and unbinding with high efficiency. However, the predefined CVs could potentially lead to certain constraints on the ligand binding pathways and conformational space. Such simulations could also suffer from the “hidden energy barrier” problem and slow convergence if important CVs were missing.^59^ Overall, the previous methods appeared computationally expensive, requiring mostly tens-of-microsecond simulations to characterize ligand binding thermodynamics and kinetics. In this context, LiGaMD2 that has allowed us to capture repetitive ligand binding and unbinding within microsecond simulations. It provides an improved approach to characterization of ligand binding thermodynamics and kinetics, especially for proteins with buried binding pockets.

## Acknowledgements

This work used supercomputing resources with allocation award TG-MCB180049 and BIO210039 through the Extreme Science and Engineering Discovery Environment (XSEDE), which is supported by National Science Foundation grant number ACI-1548562, and project M2874 through the National Energy Research Scientific Computing Center (NERSC), which is a U.S. Department of Energy Office of Science User Facility operated under Contract No. DE-AC02-05CH11231, and the Research Computing Cluster and BigJay Cluster resource funded through NSF Grant MRI-2117449 at the University of Kansas. This work was supported in part by the National Institutes of Health (R01GM132572) and National Science Foundation (2121063).

## Supporting Information

**Tables S1, S2** and **Figures S1**-**S2** are provided in the Supporting Information. This information is available free of charge via the Internet at http://pubs.acs.org.

## Table of content

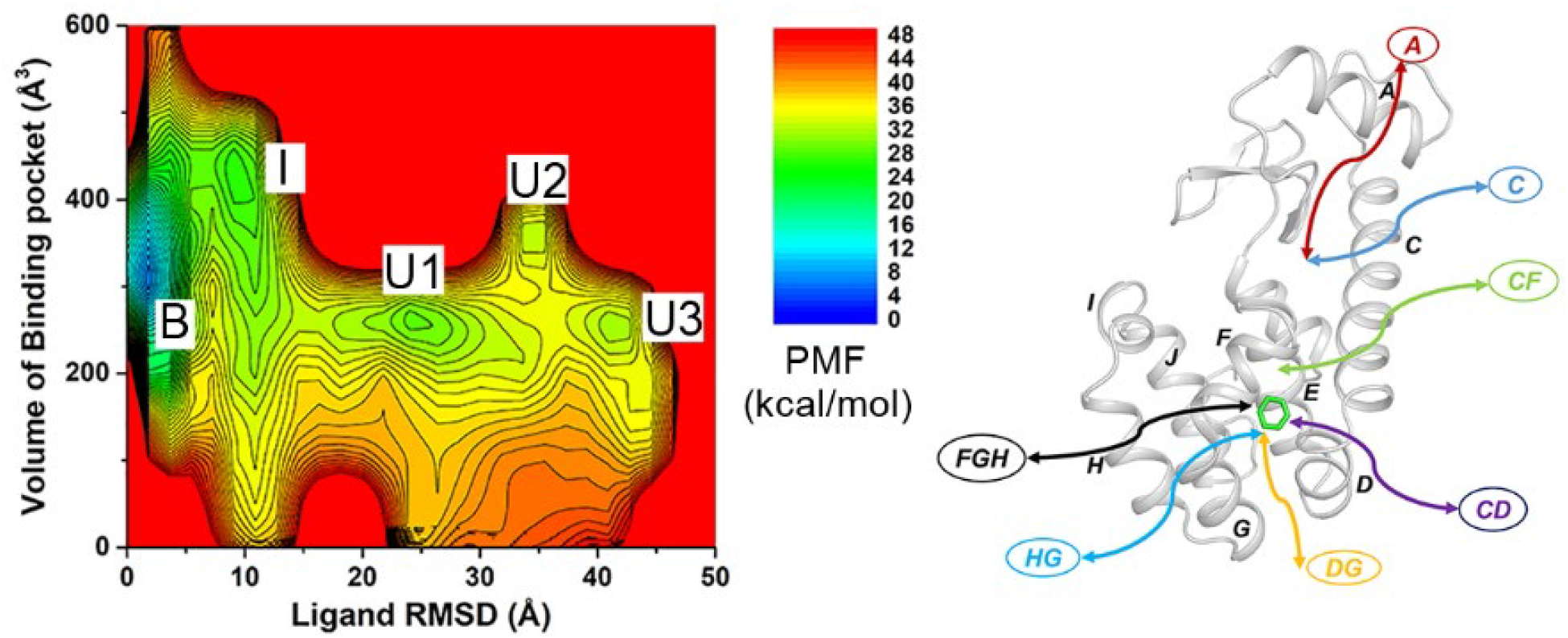

