## Supplementary material for "Ligand Gaussian accelerated molecular dynamics 2 (LiGaMD2): Improved calculations of ligand binding thermodynamics and kinetics with closed protein pocket": SI

**Table S1** The ligand bound and unbound time periods ($\tau_B$ and $\tau_U$) recorded from LiGaMD2 simulations of the ligand-T4L binding system.

| System | ID | $\tau_B$ (ns) | $\tau_U$ (ns) |
|---|---|---|---|
| L99A-BEN | Sim1 | 85.3,77.3,70.5,21.0,140.4,41.9 | 42.0,35.3,61.8,70.7,170.6,83.3 |
| | Sim2 | 212.49,26.2,19.6,13.0,111.2,32.9 | 218.80,106.3,38.0,64.6,13.2,43.8 |
| | Sim3 | 113.9,35.5,14.3,94.4,278.2 | 19.7,136.34,173.3,110.1,25.17 |
| M102A-BEN | Sim1 | 12.99,187,6,45.8,26.2 | 490.7,56.1,156.2,24.2 |
| | Sim2 | 54.94,36.0,362.6 | 240.01,281.4,25.5 |
| | Sim3 | 112.9,419.21,40.9 | 87.78,240.1,101.1 |
| F104A-BEN | Sim1 | 360.7,191.6,64.29,10.5,32.9,17.1 | 85.2,17.1,26.3,40.5,43.3,101.5 |
| | Sim2 | 456.4,47.2,18.4 | 261.1,110.6,96.3 |
| | Sim3 | 68.17,147.0,57.6,10.5,9.8,45.0,8.9 | 129.8,35.4,56.6,45.4,108.2,59.2,221.0 |
| L99A-IND | Sim1 | 6.84,26.7,14.8,100.8,20.7,9.9 | 11.85,329.0,127.5,97.8,23.0,41.5,189.9 |
| | Sim2 | 89.14,151.3,26.7 | 347.5,88.9,412.3 |
| | Sim3 | 16.87,81.4,21.0,171.7 | 284.8,73.9,217.7,122.7 |

**Table S2** Energy barriers of ligand dissociation ("off") and binding ("on") calculated from the reweighed (**Δ*F***) and modified (no reweighting, **Δ*F****) free energy profiles, curvatures of the reweighed (*w*) and modified ($w^*$) free energy profiles near the ligand Bound ("B"), Barrier ("Br") and Unbound ("U") states, and the ratio of apparent diffusion coefficients calculated from the LiGaMD2 simulations without reweighting (modified, $D^*$) and with reweighting ($D$).

| Sim | Δ*F* (kcal/mol) | | Δ*F** (kcal/mol) | | *w* | | | *w** | | | *D*/D* | |
|---|---|---|---|---|---|---|---|---|---|---|---|---|
| | Off | On | Off | On | B | Br | U | B | Br | U | Off | On |
| L99A-BEN | 10.5±0.86 | 4.93±1.38 | 2.55±0.43 | 0.57±0.14 | 9.78±9.20 | 0.95±0.29 | 0.07±0.03 | 0.87±0.81 | 0.077±0.034 | 0.02±0.001 | 4.42 | 17.67 |
| F104A-BEN | 9.00 ± 1.07 | 6.03±0.11 | 2.35 ± 0.34 | 1.15 ± 0.50 | 22.38 ± 11.42 | 2.05±0.75 | 0.15 ± 0.05 | 0.50 ± 0.11 | 0.054 ± 0.037 | 0.035 ± 0.0017 | 6.16 | 813.11 |
| M102A-BEN | 8.55 ± 0.54 | 5.40±1.07 | 2.36 ± 0.35 | 0.54 ± 0.053 | 3.11±0.29 | 1.98±1.23 | 0.57 ± 0.14 | 0.57 ± 0.14 | 0.018 ± 0.014 | 0.0028±0.0112 | 86.51 | 3519.25 |
| L99A-IND | 7.11 ± 0.34 | 4.37±0.33 | 2.23 ± 0.26 | 0.53 ± 0.10 | 2.12±0.83 | 0.14±0.09 | 0.27 ± 0.15 | 0.21 ± 0.06 | 0.14±0.07 | 0.0075±0.0017 | 26.89 | 467.56 |

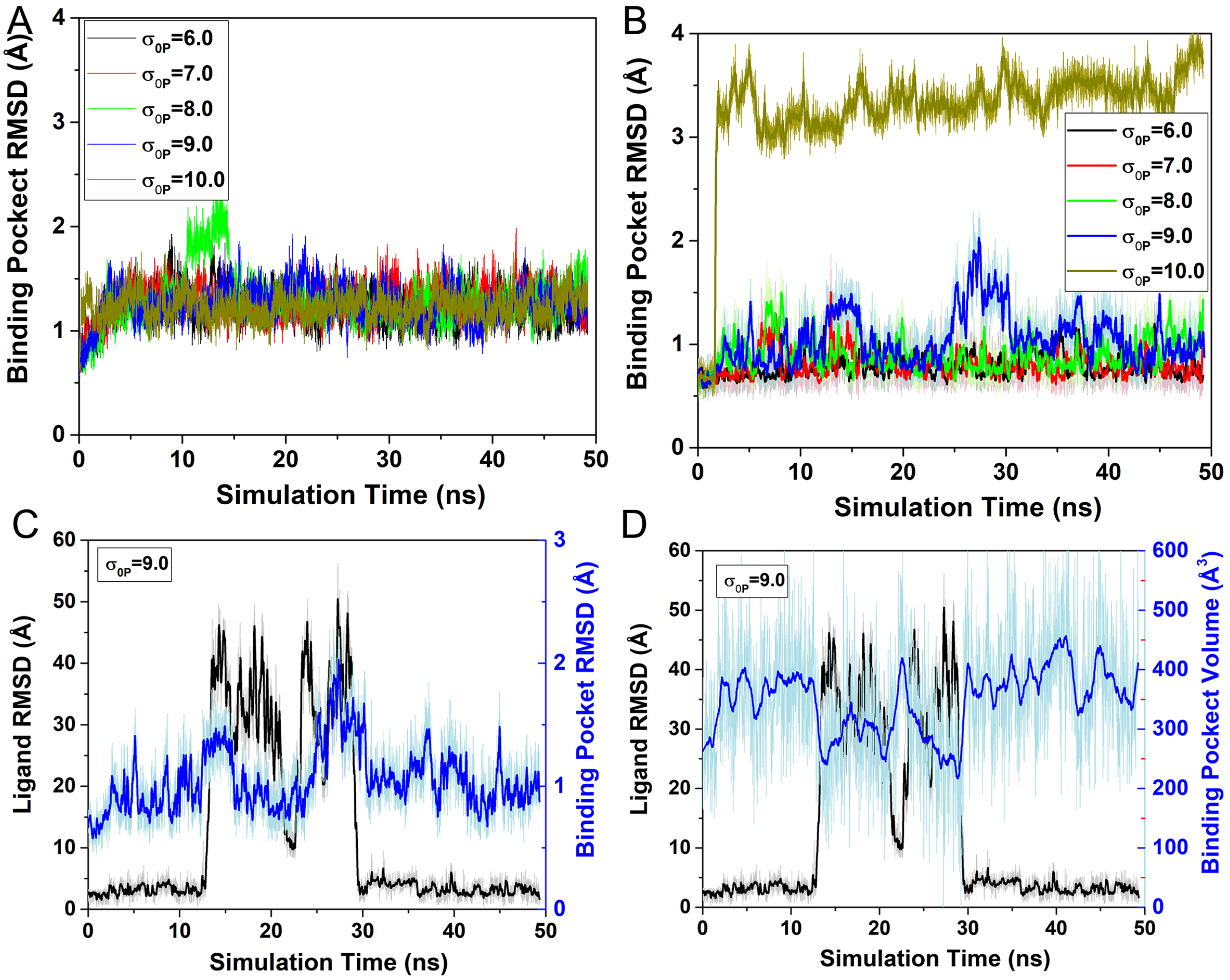


**Figure S1.** Time courses of RMSD of the ligand binding pocket relative to the X-ray structure in L99A T4L calculated from 49.2 ns LiGaMD (**A**) and LiGaMD2 (**B**) equilibration simulations; (**C**) Time courses of RMSD of the ligand relative to the X-ray structure and ligand binding pocket relative to the X-ray structure in L99A T4L calculated from 49.2 ns LiGaMD2 with $\sigma_{0P}$ at 9.0 kcal/mol. (**D**) Time courses of RMSD of ligand relative to the X-ray structure in L99A T4L and pocket volume calculated from 49.2 ns LiGaMD2 with $\sigma_{0P}$ at 9.0 kcal/mol.

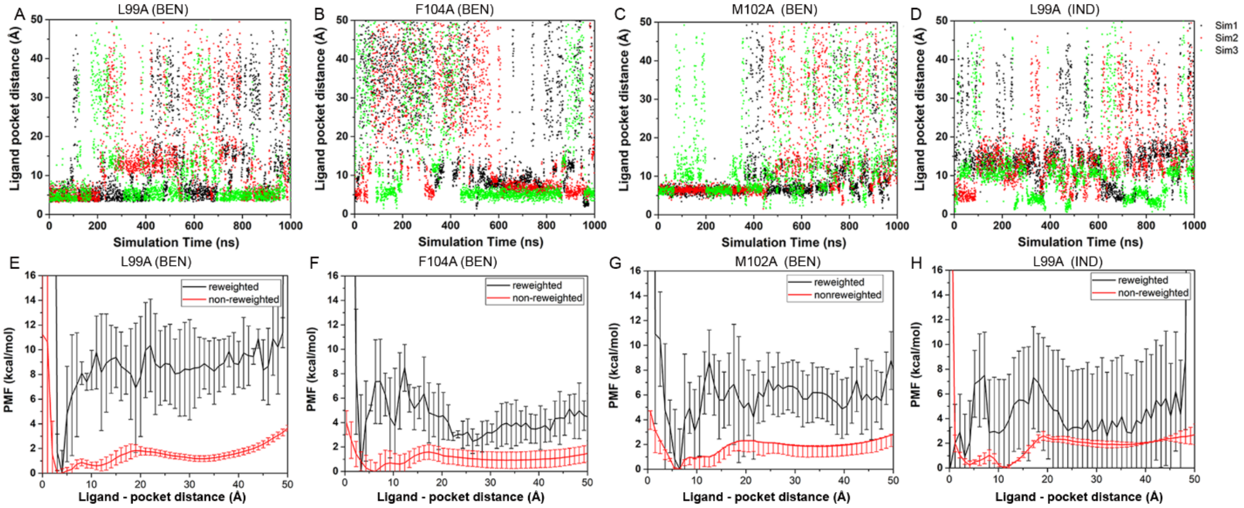


**Figure S2.** (**A**-**D**) time courses of the center-of-mass distance between the ligand and the protein pocket (defined by protein residues within 5 Å of ligand) calculated from three independent 1 μs LiGaMD2 simulations of (**A**) benzene binding to the L99A T4L, (**B**) benzene binding to the F104A T4L, (**C**) benzene binding to the M102A T4L, and (**D**) indole binding to the L99A T4L. (**E**-**H**) The corresponding PMF profiles of the ligand-pocket distance averaged over three LiGaMD2 simulations of (**E**) benzene binding to L99A T4L, (**F**) benzene binding to F104A T4L, (**G**) benzene binding to M102A T4L, and (**H**) indole binding to L99A T4L. Error bars are standard deviations of the free energy values calculated from three LiGaMD2 simulations.
